# A Tissue Microenvironment Analogous to Certain Tumor Microenvironments Facilitates HIV Persistence

**DOI:** 10.64898/2026.04.30.720299

**Authors:** Eliana U. Crentsil, Natalie Stegman, Anne Monette, Thomas J. Hope, Ramon Lorenzo-Redondo

## Abstract

The HIV reservoir that establishes early upon infection and persists in tissues remains the primary barrier to a functional cure. While progress has been made to study the reservoir in blood compartments and specific cell types, knowledge gaps remain on the tissue microenvironment that facilitates persistence. The development of a novel immunoPET/CT-guided spatial transcriptomics pipeline has enabled the localization of foci of viral infection in tissues, rare events that are challenging to sample. Prior studies leveraging the pipeline have characterized the viral microenvironment (VME) gene signatures, cell types and interactions, and host transcriptome gene drivers of viral persistence. Detailed characterization of the VME revealed multiple shared features with certain tumor microenvironments (TMEs), suggesting shared immunoregulatory and survival mechanisms. In this study, we apply the immunoPET/CT-guided spatial transcriptomics pipeline in the SIVmac239/rhesus macaque model to define immune mechanisms underlying persistent versus transient HIV/SIV reservoirs. We utilize a systematic approach to highlight correlates between the VME and TME transcriptional programs, gene pathways, local tissue neighborhoods, cell-cell interactions, and host transcriptomic drivers. Analysis of broad transcriptional programs revealed SIV-localized spatial enrichment of genes associated with multiple cancer subtypes. The persistent reservoir was characterized by gene pathways of “cold” TMEs (e.g., epithelial to mesenchymal transition, TGFβ activation) whereas the transient reservoir was comparable to “hot” TMEs with cytotoxic immune activation. Cell-cell interaction analysis identified regulatory T cells as a key mediator of interactions in both persistent and transient reservoirs. Machine learning identified KRT8, EPCAM, and RRM2, genes with known roles in mediating carcinogenesis, among the top host transcriptomic drivers of the TME phenotype of the persistent VME. Collectively, the findings of this study provide novel and transformative insights on key mechanisms of HIV/SIV persistence and reveal potential targets for immunotherapeutic strategies aimed at reservoir disruption or clearance towards a functional HIV cure.

## Introduction

Although significant progress has been made towards the treatment of HIV through advances in antiretroviral therapy (ART), the persistence of long-lived viral reservoirs remains the main barrier to achieving a functional cure as they lead to a rapid viral rebound after therapy cessation [1, 2]. To date, most HIV reservoir studies have relied on analyses of blood-derived components (e.g., peripheral blood mononuclear cells) and have focused primarily on cell types traditionally considered to harbor the majority of the reservoir, particularly CD4⁺ T cells [3, 4]. However, these blood-focused approaches miss critical aspects of viral persistence in tissues, especially in mucosal and lymphoid tissues that are likely the primary source of tissue reservoirs [3, 5, 6]. Such tissue compartments contain diverse and highly specialized immune and non-immune cell types whose local interactions, microenvironmental cues, and spatial organization can substantially influence viral persistence [4, 7]. As a result, key contributors and reservoir-supportive processes remain underexplored using conventional blood-focused methodologies [1].

To overcome this major knowledge gap, we developed an immunoPET/CT-guided spatial transcriptomics pipeline in combination with immunofluorescence (IF) viral detection, transcriptional data deconvolution, and machine learning (ML) modeling, to perform in-depth characterization of tissue viral reservoirs using the Simian Immunodeficiency Virus (SIV)mac239/rhesus macaque (RM) model [8]. This study focused on reservoirs that initiate viral rebound after analytical treatment interruption (ATI) in tissues from the gastrointestinal (GI) tract, which harbors the largest viral reservoir in the body and has been shown to be the most probable origin of the viral rebound upon ATI both in SIV and HIV-1 [1, 5]. We characterized the tissue microenvironment surrounding viral foci during the eclipse-phase of viral rebound (the earliest viral rebound phase at 4-6 days post-ATI), comparing the features of “persistent” to “transient” viral reservoirs [8].

The temporally-based reservoir categories are reflective of previous studies in the SIV/RM model which indicate that early initiation of ART (by day 4 or 5), followed by viral suppression for at least 6 months, results in a lack of viral rebound after ATI, or a delayed rebound phase that wanes within one year of treatment, which has been referred to as a “transient reservoir” [9, 10]. The transient reservoir is not a bona fide “reservoir” in the strict sense of the term with regards to long-term persistence of the virus in tissues. It is more reflective of the time required to eliminate or control all potential sources of replication-competent virus. ATI allows this extended period to be functionally defined. Unless host responses have completely eliminated all potential HIV sources, viral rebound is expected. Moreover, this phenomenon of an extended period required for varied host responses to eliminate a virus is not limited to HIV: in certain viruses such as human papillomavirus, measles, etc., viral RNA can still be detected months after the initial infection but clears within one year [11–15]. The ability to eventually control/clear SIV with early ART initiation contrasts with typical models in which ART is initiated ∼6-10 weeks after infection, leading to a persistent reservoir that is most physiologically comparable to the rapid homogeneous rebound observed in people with HIV (PWH) [9, 10, 16]. In a previous study, we defined a spatially organized viral microenvironment (VME) that governs reservoir durability and rebound potential [8]. Persistent reservoirs were localized to discrete mucosal foci and embedded within immunosuppressive, tertiary lymphoid structure–like niches dominated by regulatory T cell–centered interaction networks and enriched for innate lymphoid cells and mast cells, whereas transient reservoirs resided in immune-active environments enriched for antiviral effector cells. At the molecular level, the VME of persistent reservoirs was characterized by activation of stress-response, metabolic, mitochondrial, and cell-cycle programs coupled to repression of global cytoplasmic translation, increased cellular senescence, and engagement of the integrated stress response. Statistical modeling, machine-learning and orthogonal imaging approaches identified stress adaptation, hypoxia, cytoskeletal remodeling, and translational control as dominant predictors of viral reservoir density, supporting a model in which durable viral persistence is reinforced by a specialized, stress-adapted tissue niche [8].

Notably, several key cell and molecular features of the VME overlapped with characteristics of various tumor microenvironments (TMEs). The integrated stress response characteristic of the VME parallels the features of hypoxia, oxidative activation, and other cell stress-related responses described in several cancers [17–19]. SIV association with red blood cells, as well as the differences in viral proximity to epithelial cells versus mesenchymal cells, may reflect the angiogenesis and epithelial-to-mesenchymal transition (EMT) processes described in the TME, and, in some cases, metastatic progression [17, 20, 21]. Furthermore, contrasts in immune cell type composition of the reservoir–specifically the prevalence of immunosuppressive cells such as regulatory T cells (Tregs) versus inflammatory and cytotoxic populations such as CD8^+^ T cells–suggest that persistent and transient reservoirs share specific features that align with broader TME phenotypes, such as the “cold” and “hot” TMEs described across different cancer types. Initially, tumors were commonly described as immune-inflamed (“hot”), immune-excluded, or immune-desert (“cold”/non-inflamed) phenotypes, reflecting the abundance and spatial distribution of immune cells within the tumor microenvironment [22–25]. The “cold” TME phenotype can also extend to immunosuppressive microenvironments which are not immune deserts but are instead populated by immunosuppressive immune cells such as Tregs, tumor-associated macrophages (TAMs), and myeloid derived suppressor cells (MDSCs), blurring the distinction between different TME phenotypes [22–28]. A primary distinguishing feature between hot and cold TMEs is the high prevalence of cytotoxic CD8^+^ T cells in hot TMEs that contrasts to the immunosuppressive microenvironment of cold TMEs (i.e., recruitment of Tregs) [25, 29, 30]. Given the functional overlap between “cold” and immunosuppressed TMEs, we will refer to the immunosuppressed phenotype as the “cold” TME. Collectively, these shared features led us to hypothesize that SIV persistence in tissue reservoirs is facilitated by the establishment of a VME that recapitulates core characteristics of specific TMEs.

In the present study, we systematically characterize the features of persistent VMEs that parallel those described in various TMEs, contrasting key characteristics with the transient reservoir comparison group. Employing a systems-level spatial transcriptomics approach, we interrogate transcriptional programs, gene pathways, local tissue neighborhoods, cell-cell interactions, and host transcriptomic drivers associated with persistent SIV reservoirs. The extensive application of spatial transcriptomics in cancer research has profoundly advanced the cancer research field’s understanding of TMEs and informed the development of immunotherapy-based treatment strategies [31–33]. By leveraging the same conceptual and analytical frameworks, our findings aim to uncover targetable features of the VME that sustain viral persistence, thereby informing therapeutic strategies directed toward eradication of persistent reservoirs and, ultimately, achievement of a functional cure for HIV.

## Results

### Persistent SIV reservoirs form structured microenvironments with high expression of genes associated with various cancers

In our previous study, we identified a significant association between the VME and cancer-associated transcriptional signatures, along with signatures linked to other disease states [8]. Building on that observation, we first asked how the transcriptional landscape of persistent SIV reservoirs compares to that of various disease conditions and thus screened diseases in the enrichDO disease ontology (DO) database [34], using spatial enrichment analysis. More than 4,000 enrichDO disease pathways were evaluated with decoupleR to identify spatially enriched programs in the persistent and transient reservoirs [35]. Among the DO pathways, the most enriched pathways (adjusted p value <0.01) were those associated with cancer (**Figure 1a**), specifically pathways annotated as “organ system cancer”, “cancer”, “disease of cellular proliferation”, “endocrine gland cancer”, “liver cancer”, “hepatocellular carcinoma”, “liver carcinoma”, “cell type cancer”, “gastrointestinal system cancer”, and “carcinoma.” Notably, the high enrichment of several GI and hepatobiliary-associated cancers further supported the sensitivity of this method for resolving biologically meaningful transcriptional patterns that matched the nature of the tissues analyzed.

**Figure 1.**
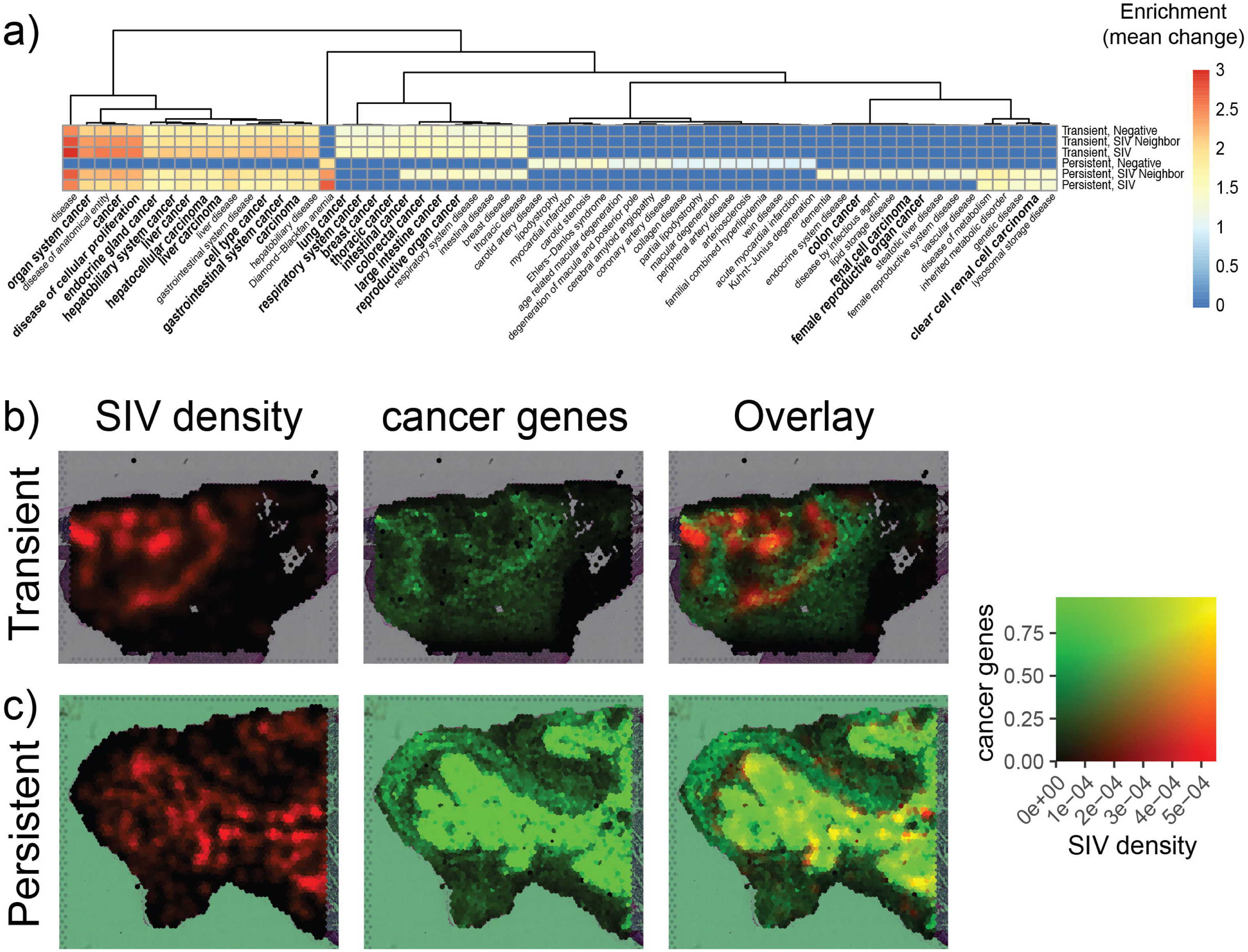
Spatial enrichment analysis of enrichDO disease pathways in spatial transcriptomics data. **A)** Relative scores of spatially enriched disease ontology pathways (adjusted p values <0.01) for the transient and persistent reservoirs by SIV region. **B)** Spatial visualization and overlay of SIV density and cancer gene pathway scores on an example slide from the transient reservoir. **C)** Spatial visualization and overlay of SIV density and cancer gene pathway scores on an example slide from the persistent reservoir. The maximum value on the color scale is the 98^th^ percentile of SIV density scores and 90^th^ percentile of the cancer gene scores for the integrated data of both the transient and persistent reservoirs (the full SIV density and cancer gene score distribution for all tissues is in **Supplementary** Figure 1).

Disease ontology scoring analysis also revealed marked differences in the spatial patterns of cancer-associated gene expression in the persistent and the transient reservoirs, as well as distinct associations with SIV foci of infection. As previously described, IF for the SIV Gag protein was used to label SIV foci of infection in an axially adjacent tissue to that used for spatial transcriptomics (separated by a distance of 20μm), classifying spots into three categories: SIV-positive, SIV-neighbor, and SIV-negative [8]. When spatially visualizing the cancer-associated gene pathway scores and the SIV density distributions individually and overlaid onto each tissue slide, the persistent reservoir had enrichment of cancer-associated gene pathways that was highly localized to the SIV-positive tissue regions, while there was comparatively low or near-absent expression in the SIV-negative tissue regions (**Figure 1c, Supplementary Figure 1**). SIV-negative regions, in turn, had relatively higher expression of gene pathways of extracellular matrix, collagen, and fibrotic-associated processes, such as “collagen disease” and “Ehlers-Danlos syndrome.” In the transient reservoir, the cancer gene spatial distribution was similarly localized to tissue regions with higher SIV density; however, the spatial signal for collagen and fibrotic-associated processes was not significant in the transient reservoir (**Figure 1b, Supplementary Figure 1**).

Together, these findings suggest that persistent reservoirs are established within spatially organized tissue niches characterized by local activation of cancer-associated transcriptional programs in SIV-associated and adjacent regions embedded within a surrounding fibrotic environment.

### Tumor microenvironment scoring reveals differential distribution of immune and stromal transcriptional features in persistent compared to transient SIV reservoirs

The disease ontology enrichment analysis, in conjunction with our previous studies on biomarkers and features of the persistent VME, indicated that persistent SIV reservoirs share transcriptional features with various cancers. While the TME is highly variable among different cancer subtypes, it is broadly classified using three components: immune, stromal, and extracellular matrix (ECM) [36, 37]. Consequently, our next step was to further characterize the shared features of the VME and TME by understanding the distribution of the individual TME components across the persistent and transient reservoirs.

To understand these patterns, we applied the Estimation of STromal and Immune Cells in MAlignant Tumor tissues using Expression data (ESTIMATE) TME scoring method [38]. This method provides “immune” infiltration scores determined by the transcriptional levels of genes associated with the amount of immune cell infiltration into tumors, and “stromal” scores that reflect the extent of stromal infiltration. The sum of these two scores is captured by the ESTIMATE score and represents the overall tumor purity (**Supplementary Figure 2**) [38]. Overall, ESTIMATE scoring recapitulated previously detected transcriptional patterns and tissue structures observed with imaging methods. For example, ESTIMATE immune scoring showed robust co-localization with a previously identified immune aggregate in a transient reservoir slide (shown in the yellow and black boxes in **Supplementary Figure 3a** and 3b, respectively) [8]. Therefore, we proceeded to analyze the associations between the spatial distribution of SIV Gag detection (**Figure 2a**) and the ESTIMATE, immune, and stromal scores (**Figure 2b**).

**Figure 2.**
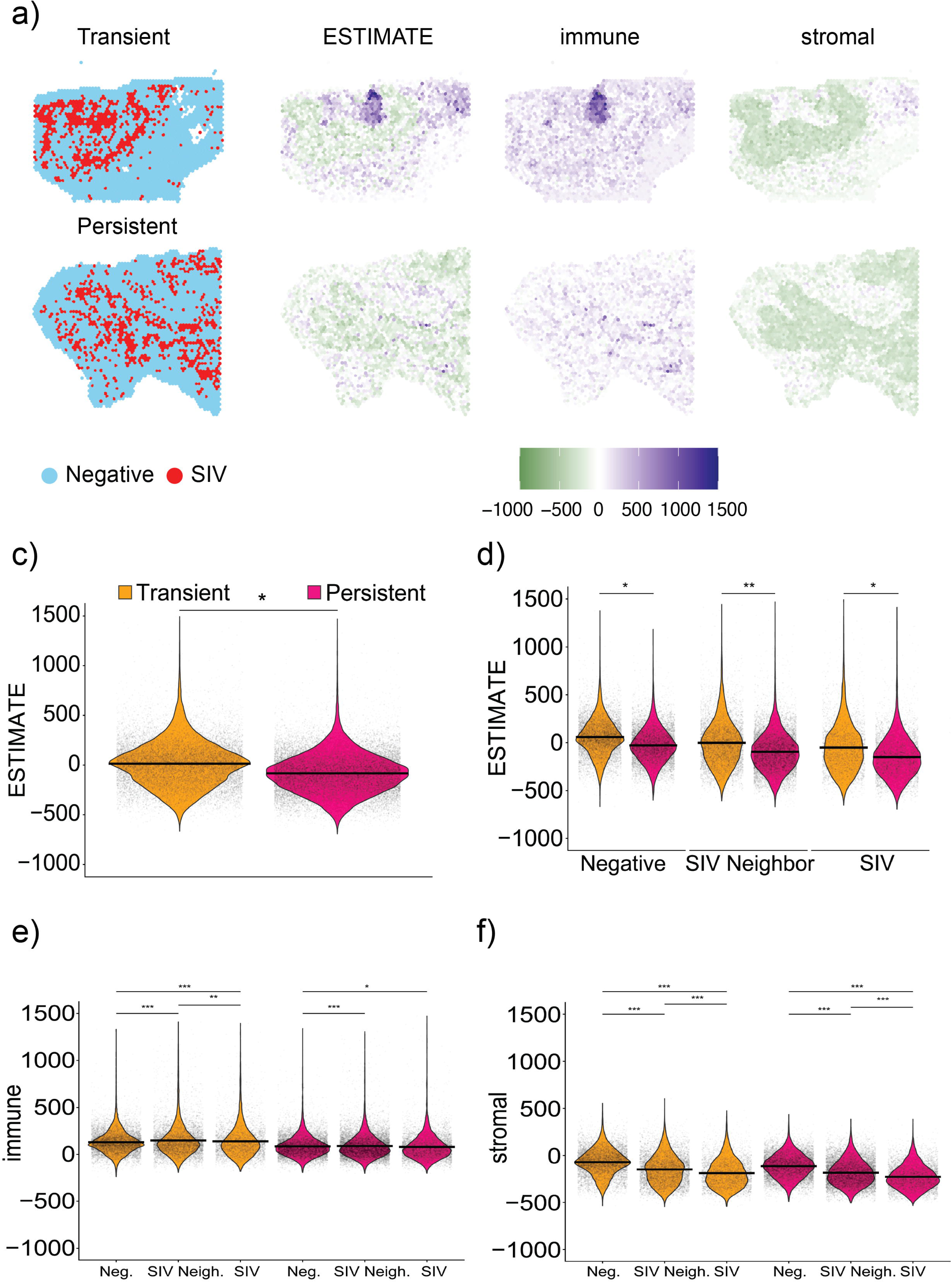
ESTIMATE TME scoring. **A)** SIV labels visualized for all spots on an example slide from the transient and persistent reservoir. **B)** ESTIMATE, stromal, and immune scores on the example slide from the transient and persistent reservoirs. **C)** ESTIMATE score numerical distribution for transient vs. persistent reservoirs. **D)** ESTIMATE score numerical distribution for transient vs. persistent reservoirs, stratified by presence of SIV foci of infection. **E)** Immune and **F)** stromal score numerical distributions for transient vs. persistent reservoirs, stratified by presence of SIV foci of infection.

**Figure 3.**
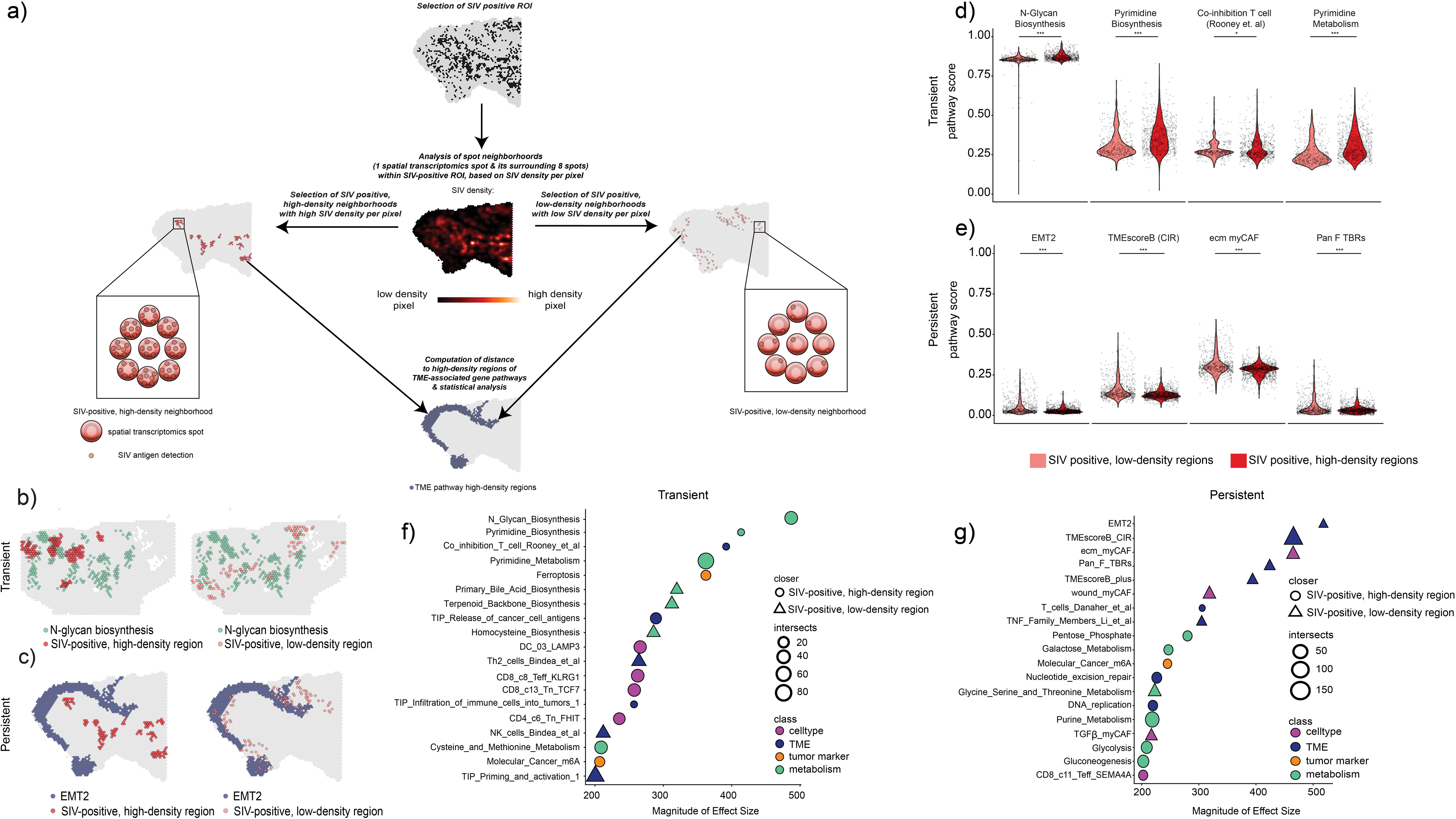
Neighborhood analysis reveals different local microenvironments in transient and persistent reservoirs. **A)** Schematic of method of geostatistical comparison of TME pathway proximity to SIV-positive, high-density and SIV-positive, low-density regions. Example image of top statistically significant enriched pathway in the **(B)** transient and **(C)** persistent reservoirs. Violin plot visualizing scores of top enriched pathways in SIV-positive, high-density and SIV-positive, low-density regions in the **(D)** transient and **(E)** persistent reservoirs. Dotplot of top enriched TME pathways in **(F)** transient and **(G)** persistent reservoirs based on mixed linear effects modeling statistical analysis of minimum Euclidean distances between TME pathway high-density regions and SIV-positive high- or low-density regions.

Mixed linear effects modeling showed that the persistent reservoir had significantly lower ESTIMATE scores than the transient reservoir, indicative of a more TME-like transcriptional status (p <0.05) (**Figure 2c**). This pattern remained consistent after stratification by the presence of SIV foci of infection (**Figure 2d**). The observed spatial distribution of ESTIMATE scores also supported the association of SIV presence with reduced composite ESTIMATE scores, which is correlated with a more TME-like phenotype. Similar trends were observed in pseudobulk analyses, indicating that these patterns were not limited to spot-level inference (**Supplementary Figure 4**).

With regards to immune infiltration specifically, the persistent reservoir displayed significantly increased immune scores when comparing SIV-negative spots to either SIV-positive or SIV-neighbor spots, while no differences were observed between SIV-positive and SIV-neighbor spots. The differences in immune scores suggest that there is limited immune infiltration into the SIV-associated regions in the persistent reservoir (**Figure 2e**). By contrast, the transient reservoir showed statistically significant stepwise increases in the immune infiltration scores from the SIV-negative to SIV-neighbor to SIV-positive spots, indicative of widespread immune infiltration. Stromal infiltration scores were significantly higher in the SIV-negative spots compared to the SIV-positive and SIV-neighbor spots in both the persistent and transient reservoirs, concordant with our previously reported negative association of SIV foci with mesenchymal cells and the stromal/fibrotic gene enrichment in SIV-negative regions identified in the spatial enrichment analysis **(Figure 1a)** [8].

In summary, this quantitative ESTIMATE scoring indicated that persistent reservoirs are embedded within a TME-like microenvironment marked by limited immune infiltration while transient reservoirs displayed widespread immune presence associated with SIV detection.

### Local tissue neighborhood analyses reveal that the persistent reservoir shares similarities to “cold” TMEs while the transient reservoir shares similarities to “hot” TMEs

The previous assessments of features present throughout the tissue samples prompted deeper investigations into shared hallmarks at the local tissue neighborhood, cell, and molecular level of the VME that reflect the TME. Thus, we next sought to understand specific hallmarks of the local microenvironment of SIV foci of infection that reflect the TME. With this goal, we used geostatistical analysis with SpottedPy [39] to identify TME-associated pathways that were in close spatial proximity to SIV foci. Analyses were completed via the selection of regions of interest (ROIs) on each slide specific to the presence of SIV foci of infection. SIV infected cell density – which we will refer to as “SIV density” – per 10x Visium spot was determined by quantifying Gag antigen detection density from IF on an adjacent section separated by 20μm, as previously described [8]. Within the ROI containing SIV-positive spots, we analyzed local neighborhoods comprising one Visium spot and its eight surrounding spots, and classified them as “SIV-positive, high-density” or “SIV-positive, low-density” based on whether local SIV density exceeded or fell below that expected under spatial randomness (**Figure 3a**).

Upon geostatistical identification of these SIV-positive, high-density regions and SIV-positive, low-density regions, we sought to identify how specific gene pathway hallmarks of TMEs are spatially distributed with respect to the SIV regions. To this aim, we compared the SIV geostatistical distributions to the expression of the gene pathways in the ImmunoOncology Biological Research (IOBR) Database. IOBR currently contains over 320 gene pathways categorized into four main classes: TME, tumor marker, cell type, and metabolism [40]. We determined neighborhood regions of high scores of the IOBR TME-associated gene pathways using the same geostatistical analysis and examined their distribution within SIV-positive, high-density regions and low-density regions in both animal groups. This analysis revealed marked differences between persistent and transient reservoirs in the pathways positioned nearest to SIV-positive high-density versus low-density regions, suggesting differences in the structures and genetic signatures within localized neighborhoods of the reservoirs. In the persistent reservoir, many of the top pathways with the largest effect size were closer to the SIV-positive, low-density regions, with notable pathways including epithelial to mesenchymal transition (EMT2), TMEscoreB_CIR (a stromal-type TME score previously identified through prognostic scoring), pan-fibroblast transforming growth factor beta (TGFβ) response signature (Pan_F_TBRS), cancer-associated fibroblasts (ecm_myCAF and wound_myCAF [extracellular matrix-producing and wound myofibroblastic cancer associated fibroblasts]) and Tumor Necrosis Factor (TNF) Family Members (**Figure 3c,g and Supplementary Figure 5b**). Collectively, most of these pathways were characteristic of stromal and fibrotic processes that tend to be associated with “cold” TME signatures [36]. In addition, a number of pathways were closer to the SIV-positive, high-density regions; these were mostly associated with metabolic pathways (e.g., pentose phosphate; galactose metabolism; glycine, serine, and threonine metabolism; purine metabolism, glycolysis, and gluconeogenesis) and DNA replication and repair (e.g., nucleotide excision repair) processes, with limited association with T cell processes. The distinct transcriptional programs observed in SIV-positive high-density versus low-density regions suggest the presence of two biologically distinct SIV-infected cell populations. This finding is consistent with our prior identification of two SIV-associated transcriptional clusters, which also corresponded to two phenotypically distinct SIV-infected cell populations detected by IF [8]. One population likely represents long-lived viral reservoirs, whereas the other appears to correspond to newly expanding viral infection.

In contrast, the pathways with the highest effect sizes in the transient reservoir tended to be closer to SIV-positive, high-density regions (**Figure 3b, f and Supplementary Figure 5a)**, including metabolic pathways (N-Glycan biosynthesis, pyrimidine metabolism and biosynthesis), Co-inhibition of T cells, CD8^+^ T cell-associated processes, as well as release of cancer cell antigens and immune infiltration (“tracking tumor immunophenotype [TIP] Release of cancer cell antigens” and “TIP Infiltration of immune cells into tumors”), consistent with the immune infiltration observed in the ESTIMATE scoring (**Figure 2e**). As opposed to the persistent reservoir, these pathways tend to be associated with the “hot” TME phenotype [30, 36]. Other notable pathways proximal to the SIV-positive, high-density regions included various metabolic processes (e.g., primary bile acid, terpenoid backbone, and homocysteine biosynthesis) and other immune cell types (e.g., Th2 cells, NK cells).

Additionally, when analyzing the broader categories of the pathways with the highest effect sizes, the most frequent category in the persistent reservoir was from the “TME” class of the IOBR database (**Figure 3g**), compared to the transient reservoir in which top pathways tended to be associated most frequently with metabolic processes, in addition to TME and immune cell type-specific processes (**Figure 3f**). These differences suggested that persistent reservoirs are associated with broader microenvironment changes whereas transient reservoirs are rather associated with changes that occur at the cell and molecular level. Additionally, when comparing the top significant pathways in **Figure 3f,g** that were unique to the transient and persistent reservoirs, “TIP Infiltration of immune cells into tumors_1” was significantly enriched in the transient reservoir, but not the persistent reservoir, which is consistent with our previous ESTIMATE score findings (**Figure 2, Supplementary Figure 5c-f**).

Finally, we also sought to determine the relative expression of the pathways within the SIV-positive high- and low-density regions. Here, mixed linear effects modeling was completed within the high-and low-density spots to compare the gene pathway scores for the top 5 pathways of the persistent and transient reservoirs. As shown in **Figure 3d-e**, the pathways that were closer to SIV-positive, high-density regions had significantly higher gene pathway expression scores in the high-density regions compared to the low-density regions. Conversely, pathways closer to SIV-positive, low-density regions had significantly higher scores in low-density regions compared to high-density regions. The corresponding expression trends within the local neighborhoods supports the specificity of the pathway expression to the high- or low-density regions of the tissues and is indicative of both local and broad tissue microenvironment remodeling.

Overall, these data indicate that both persistent and transient reservoirs exhibit TME-like features, but that the local transcriptional programs surrounding SIV foci differ fundamentally between them. Persistent reservoirs reflected “cold” TMEs, while the signatures in transient reservoirs were analogous to “hot” TMEs.

### Regulatory T cells have central, but mechanistically different, roles in the microenvironment of persistent and transient reservoirs

To delineate the shared hallmarks of the VME and TME at the cellular level, we next analyzed cell-cell interactions to obtain a more granular and dynamic assessment of the immune microenvironment in persistent and transient reservoirs. To this end, spatially-based cell-cell interactions were identified using Spatial Omics Analysis in Python (SOAPy) [41]. Interactions were inferred at both short range (contact, adjacent spatial transcriptomics spots) and long range (secretory, non-adjacent spots) within tissues. Immune cell types and frequencies at each 10x Visium spatial transcriptomics spot were deconvoluted and contact and secretory interactions were determined based on the cell type label of the maximum frequency immune cell type at each spot [8]. In order to provide additional detail on subtypes of cells interacting in the VME, cell type labels were merged with the SIV-related spot labels (SIV-positive, SIV-neighbor, and SIV-negative) [8]. Focused analysis on immune cell interactions revealed a high involvement of regulatory T cells (Tregs) in the interactions comprising both the persistent and transient reservoirs; however, the types of interactions in which Tregs engaged differed between the reservoirs (**Figure 4a, Supplementary Figure 6**).

**Figure 4.**
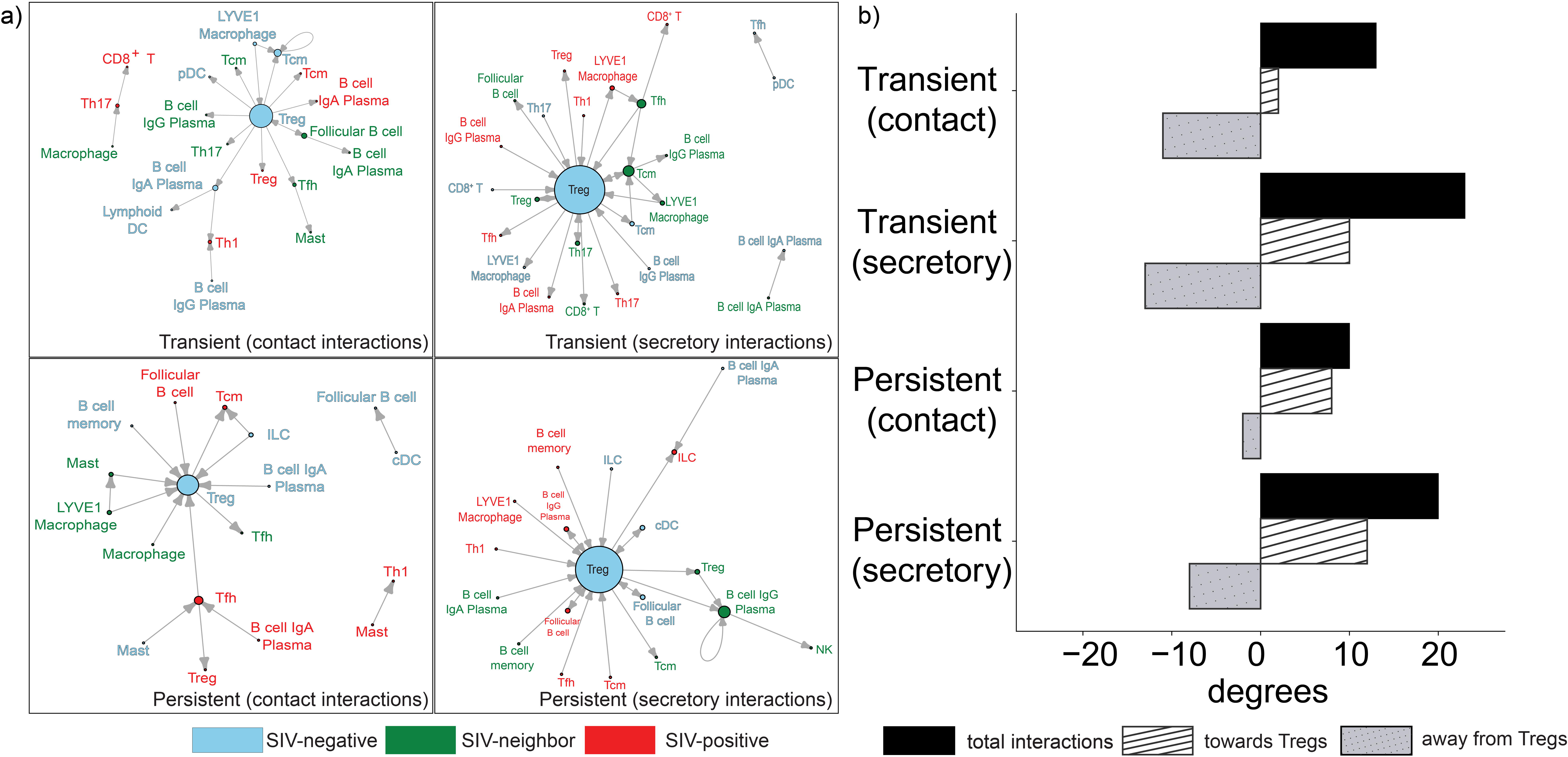
Short- and Long-range cell-cell interactions in the viral microenvironment (VME) **A)** Short-range (contact) and long-range (secretory) immune cell-cell interactions of the VME for the transient and persistent reservoirs. **B)** Degrees of centrality in cell interaction network for Tregs in the transient and persistent reservoirs, including the total number of cell interactions that involve Tregs as well as stratification into the number of interactions toward and away from Tregs.

In the persistent reservoir, SIV-negative Tregs predominantly received rather than transmitted signals (**Figure 4b, Supplementary Figure 7**). Among the interactions received by SIV-negative Tregs, the majority of the short-range, contact interactions involved innate immune cells, including mast cells, ILCs, and macrophages, whereas long-range, secretory interactions involved a broad variety of cell types, with the highest counts of interactions being received from memory B cells and LYVE1 macrophages. In the transient reservoir, SIV-negative Tregs were also involved in the highest counts of cell-cell interactions but sent out signals to other immune cells rather than receiving them (**Figure 4a**). Approximately half of the contact interaction signals sent from Tregs in the transient reservoir were to adaptive immune cells: Tcm, Th17, Tfh. Notably, Th17 cells in SIV-positive spots were involved in contact interactions with CD8^+^ T cells also in SIV-positive spots, possibly leading to cytotoxic immune activation akin to the “hot” TME phenotype. Similarly, for the secretory interactions, the majority of signals sent out by SIV-negative Tregs were also towards adaptive immune cells, including CD8^+^ T cells, Tcm, Tfh, and Th17. Overall, innate immune cells signaling to Tregs in the persistent reservoir may mediate the immunosuppressive microenvironment associated with “cold” TMEs while the signals being sent from Tregs to adaptive immune cells in the transient reservoir may drive its immune-active, “hot” TME phenotype [31, 37, 43].

In summary, these cell-level analyses revealed that Tregs play a crucial role in the VME, as has previously been reported for the TME [42–44]. However, the directionality and cell types involved in the interactions with Tregs differ in the persistent versus transient reservoirs: generally, Tregs received signals from innate cell types in the persistent reservoir, while Tregs sent signals to adaptive immune cells in the transient reservoir. Notably, interactions were present across all SIV regions (SIV-positive, SIV-neighbor, SIV-negative), demonstrating that cell-cell signaling in the reservoir may be mediated by molecular processes occurring throughout the tissue microenvironment.

### Host transcriptome genes that drive TME phenotypes and may mediate reservoir persistence

Next, we sought to dig deeper into specific genetic drivers of the TME score and used machine learning to provide mechanistic insights on the TME-like phenotype and to identify biomarkers that can serve as potential therapeutic targets of the reservoir [45]. We applied Extreme Gradient Boosting (XG Boost) regression to spatial transcriptomic profiles from persistent and transient reservoirs and ranked genes by their contribution to prediction of the ESTIMATE TME score. The top five TME score drivers unique to the persistent reservoir revealed by the machine learning models were keratin 8 [KRT8 (ENSMMUG00062792)], epithelial cell adhesion molecule [EPCAM], malectin [MLEC], ribonucleotide reductase regulatory subunit M2 [RRM2], and carbonic anhydrase 1 [CA1] **(Figure 5b, d)**. Conversely, keratin 18 [KRT18 (ENSMMUG00000031911)], tetraspanin 8 [TSPAN8], SAM and SH3 Domain Containing 3 [SASH3], chloride channel accessory 1 [CLCA1], and ATPase Na+/K+ transporting Subunit Alpha 1 [ATP1A1] were the top five gene drivers of the TME score unique to the transient reservoir (**Figure 5a, c**).

**Figure 5.**
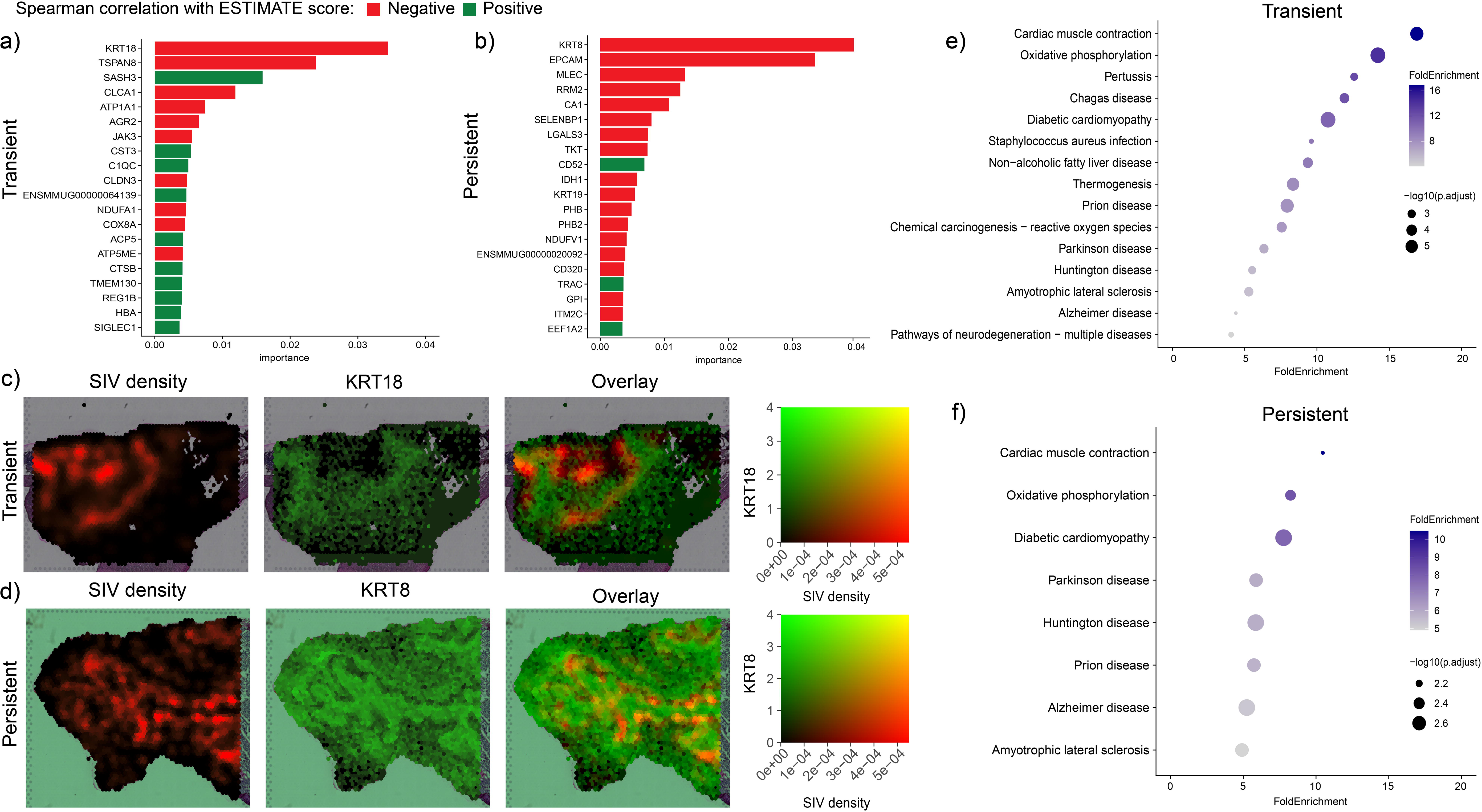
Identification of gene drivers of the TME-like phenotype. Top gene drivers identified via machine learning that are unique to the **(A)** transient and **(B)** persistent reservoirs, ranked by importance score. The color of the bars indicates whether the spearman correlation between the gene expression (SC Transform data) and the ESTIMATE score is negative or positive. Lower ESTIMATE scores are associated with the TME phenotype, so a negative correlation suggests that higher host transcriptome gene expression may be associated with a more TME-like phenotype. KEGG pathway enrichment analysis of the unique gene drivers among the top 100 ranked by importance for the **(C)** transient and **(D)** persistent reservoirs. Spatial visualization of SIV density, expression of the top ranked gene driver, and overlay for the **(E)** transient and **(F)** persistent reservoirs.

Several of these top gene drivers identified by the machine learning models have been linked to the TME and carcinogenic processes. KRT8 was the top gene driver unique to the persistent reservoir, whereas KRT18 was the top gene driver in the transient reservoir (**Figure 5a,b**). KRT8/18 are co-expressed intermediate filaments of epithelial cells that play a key role in providing support for cellular structures. KRT8/18 have been identified as diagnostic markers in certain cancers, and their loss has been associated with EMT and cancer metastasis, which was among the pathways upregulated in the neighborhood analysis in the persistent reservoir (**Figure 3c, g**) [46, 47].

The second-ranked persistent reservoir TME score driver was EPCAM, a well-established diagnostic marker for various epithelial cancers [48, 49]. In contrast, the second-ranked TME score driver in the transient reservoir, TSPAN8, has been associated with gastric cancer tumorigenic processes (e.g., proliferation, invasion) and, like the keratins, has roles in aspects of cell surface structure and signaling [50–52].

To extend these findings from individual genes to pathway-level mechanisms, we performed KEGG enrichment on unique TME score gene drivers among the top 100 ranked by importance in the persistent and transient reservoirs [53]. In the persistent reservoir, top TME score drivers enriched for pathways associated with oxidative phosphorylation and a number of neurodegenerative conditions that have been linked to the integrated stress response, endoplasmic reticulum stress, among other oxidative processes (e.g., Alzheimer’s, Huntington’s, and Parkinson’s disease) (**Figure 5f**) [54–56]. A similar pattern was noted among the top TME score drivers unique to the transient reservoir, although to a lesser degree (**Figure 5e**). The enriched KEGG pathways are consistent with our previous findings on the role of oxidative and integrated stress pathways in persistent reservoir establishment [8].

### Multiple top drivers of the TME phenotype in the persistent reservoir have been reported to be upregulated in human tumor samples

After identifying gene drivers associated with the TME score in both the persistent and transient reservoirs using machine learning, we next explored which of these genes may be the most translationally relevant based on their expression in human tumor samples. To this end, we explored publicly available data from The Caner Genome Atlas (TCGA) to identify which of the top drivers of the TME score in persistent and transient reservoirs have been implicated in human cancers [57]. Among the top five gene drivers of the TME score in the transient reservoir, multiple had relatively higher frequencies of mutations across various cancer subtypes, including skin and uterine cancers, compared to other cancer types in the TCGA (**Supplementary Figure 8a**). However, in the persistent reservoir, uterine corpus endometrial carcinoma (UCEC) was consistently the most frequent cancer type to have mutations in each of the five top gene drivers of the TME score (**Supplementary Figure 8b**). All top five gene drivers of the TME score in the persistent reservoir being associated with UCEC suggests shared transcriptional programs between the VME and the TME of UCEC. Some of the hits of interest with pertinent roles in carcinogenesis among the top drivers found in the persistent reservoir are indicated in **Figure 6a**. Of note, this association of UCEC with transcriptional features of the TME score of the persistent reservoir also appears consistent with the disease ontology analysis in **Figure 1** in which “female reproductive organ cancer” and “female reproductive system disease” were both spatially enriched specifically in the persistent reservoir.

**Figure 6.**
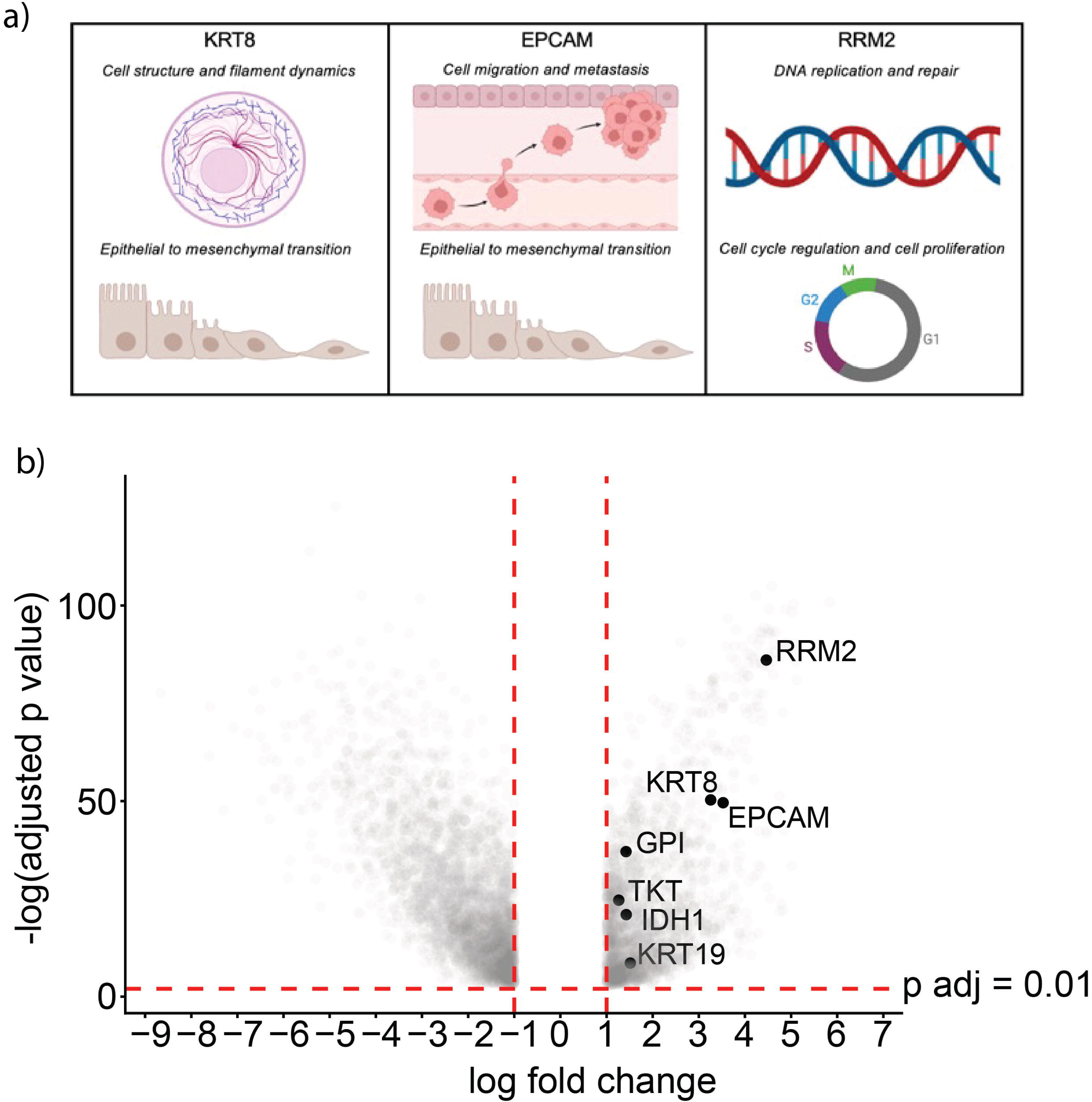
Top gene drivers of the persistent reservoir are upregulated in uterine corpus endometrial carcinoma (UCEC). **A)** Key cellular functions of top gene drivers of the TME-phenotype in the persistent reservoir. (Created with BioRender.com) **B**) Top gene drivers of the TME-phenotype in the persistent reservoir were statistically significantly upregulated in RNA sequencing data published in Luo et.al. comparing UCEC tissues to control endometrial tissues (data values shown are from Luo et. al. RNA sequencing data).

Consequently, we sought to determine whether these top gene drivers of the TME score in the persistent reservoir are differentially expressed in UCEC towards further delineating which biomarkers may have clinical relevance for potential translational applications. In assessing these top drivers (**Figure 5b)** in the context of RNA-sequencing data comparing UCEC to normal endometrial tissues from the TCGA and University of California, Santa Cruz (UCSC) Xena databases (Luo et. al.), we found 7 among the top 20 unique gene drivers were statistically significantly upregulated in UCEC (**Figure 6b**) [58]. In addition to KRT18 and EPCAM, which both have known roles in carcinogenesis, the gene hit that was the most significantly upregulated, RRM2, has also been implicated in carcinogenic processes, primarily through its roles in DNA replication and repair as well as cell cycle regulation and cell proliferation (**Figure 6a**) [59]. Consequently, RRM2, KRT8, EPCAM, GPI, TKT, IDH1, and KRT19 may be targets of interest for translational applications in leveraging immunotherapeutic approaches that have been used to target UCEC towards targeting persistent HIV/SIV reservoirs.

## Discussion

In this study, we leveraged spatial transcriptomics across tissues harboring persistent and transient SIV reservoirs to further define the VME at systems-level resolution. Our findings reveal that persistent reservoirs adopt transcriptional, cellular, and architectural features strikingly analogous to immunologically “cold” TMEs, whereas transient reservoirs resemble “hot,” immune-infiltrated TMEs. This parallel extends beyond descriptive similarity, suggesting that common microenvironmental principles govern both tumor immune evasion and HIV/SIV persistence.

Our analysis showed that SIV-associated tissue regions were spatially enriched for cancer-related gene signatures in both the persistent and transient reservoirs, with spatial localization of these signatures near regions with high density of viral detection. This convergence was evident when comparing shared transcriptional features between specific components of the TME (e.g., immune and stromal) within the reservoirs. In persistent reservoirs, stromal infiltration was lower in SIV-positive areas whereas immune infiltration was comparable between SIV-positive and SIV-neighbor regions. This result indicates that persistence is not driven by simple immune exclusion, but rather by the establishment of a locally organized, immunosuppressed niche. In contrast, transient reservoirs showed similarly low stromal infiltration; however, they exhibited pronounced immune infiltration within SIV-positive regions, consistent with effective immune surveillance of these foci.

Neighborhood-level geostatistical analysis provided detailed exploration of the microenvironment surrounding SIV-associated regions. In persistent reservoirs, stromal elements were relatively depleted in the immediate vicinity of SIV high-density regions, yet enriched in surrounding low-density viral regions, where programs associated with epithelial–mesenchymal transition (EMT), cancer-associated fibroblasts, and extracellular matrix remodeling were prominent. The localization of these programs around, rather than within, SIV-dense regions suggests the formation of protective structures that “wall off” infected cells from immune clearance, analogous to the immune evasion observed in “cold” tumors resistant to immunotherapy. In contrast, transient reservoirs displayed marked immune infiltration and activation of cellular processes –specifically related to cytotoxic T cell processes– near SIV-positive, high-density regions, indicative of immune-active niches resembling “hot” TMEs. Additionally, metabolic processes were activated in both the persistent and transient reservoirs, which may be indicative of the shunting of metabolites towards the VME, much like the metabolic dysregulation characteristic of tumor cells and the TME. Accordingly, there are emerging cancer therapeutic approaches that incorporate metabolic targeting in conjunction with immunotherapy which may also be valuable considerations for persistent reservoir clearance [30, 60]. Our previous findings indicate the presence of two populations of rebounding foci of infection during the eclipse phase of rebound, consistent with the expected newly infected cells that are expanding and disseminating throughout the tissue after ATI, and the stable reservoir population that allowed the virus to survive during ART. This was also supported by IF validation in which we found two SIV-infected populations: a population of cells that were SIV Gag-positive and positive for phosphorylated eIF2α (p-eIF2α), a key mediator of the shutdown of cap-dependent translation in the integrated stress response (ISR), and another that is SIV Gag-positive and p-eIF2α negative [8, 61]. The IF validated our spatial transcriptomics pipeline’s sensitivity to detect the two expected populations, and confirmed the significant association between SIV reservoirs and activation of the ISR. Based on the local tissue neighborhood analysis performed in this study, we postulate that the SIV Gag-positive, high-density regions with metabolic and immune activation (CD8^+^, NK, Th2 cells), are associated with the newly infected population, whereas the SIV Gag-positive, low-density regions are the cells maintaining the persistent reservoir, characterized by broad microenvironment-associated remodeling (e.g., EMT) [8]. To expand the analysis of these two populations, we are currently developing imaging and transcriptional methods to perform detailed characterization of these populations at the single cell level.

Deeper investigation of the cellular interactions driving the TME-like phenotypes revealed that Tregs are key modulators in both the persistent and transient reservoir microenvironments. In persistent reservoirs, Tregs were inferred to receive signals from innate immune cells, reinforcing the link with an immune-excluded VME. Moreover, previous spatial analysis of inferred Treg frequencies showed increased Treg presence in SIV-dense regions of persistent reservoirs, comparable to the immunosuppressive cell types often present in “cold” tumors [8, 36, 62]. In transient reservoirs, however, Tregs sent signals out to engage various adaptive immune cells in the microenvironment, consistent with immune-active regions. The prominent signaling axis from Th17 to cytotoxic CD8^+^ T cells observed in transient reservoirs –condition in which animals were infected through intravaginal or intrarectal challenge– aligned with the early targeting of Th17 cells in the mucosa upon HIV/SIV mucosal infection and their role in orchestrating downstream immune activation. [8, 63, 64]

Gene driver analysis of TME-like programs revealed distinct molecular states underlying persistent and transient reservoirs. Among the top gene drivers for TME signatures in persistent reservoirs, we found genes that enriched for epithelial and proliferative pathways, including KRT8, EPCAM, MLEC, RRM2, and CA1, consistent with a tumor cell–intrinsic program linked to structure, adhesion, and growth [65, 66]. Notably, EPCAM is a clinically targeted tumor antigen [48, 49, 67–70], while RRM2 and ATP1A1 represent host-dependency pathways with known links to viral replication and immune modulation [71–73]. In contrast, transient reservoirs were characterized by KRT18, TSPAN8, SASH3, CLCA1, and ATP1A1, suggesting a more heterogeneous epithelial–immune interface with features of immune activation and differentiation [74, 75]. Several of these drivers have established roles in tumor biology and therapeutic response. The association of some of these transient reservoir TME drivers (e.g., CLCA1, ATP1A1) with relatively higher mutation frequencies in “exceptional responder” phenotypes –i.e., patients who respond to treatments and have significantly longer survival times than others with a clinically comparable condition–[76], further supports a link between transient reservoir programs and heightened immune responsiveness. Consistently, oxidative phosphorylation and stress-response pathways were enriched across both reservoir types, aligning with the metabolically active VME previously reported [8]. Finally, comparisons with publicly available cancer datasets revealed that some of the identified persistent reservoir gene drivers are upregulated in human cancers, including uterine corpus endometrial carcinoma, suggesting conserved microenvironmental architectures between the VME associated with persistent reservoirs and specific TMEs.

The VME features and potential biomarkers identified in this study reveal key insights and directions for immunotherapeutic approaches to targeting the persistent HIV reservoir. Modulation of EMT, TGFβ, and genes associated with stromal activation and the resulting ECM deposition may enable persistent tissue reservoir targeting for eradication [77, 78]. TGFβ, in particular, has been previously implicated in both HIV pathophysiology and “cold” TME phenotypes [20, 79–85]. The enrichment of such pathways in the SIV-positive, low-density regions suggests local microenvironment remodeling that extends beyond the high-density regions of SIV foci of infection specifically. Inhibition of immunosuppressive signaling in persistent reservoirs can be coupled with stimulating cytotoxic immune cell activity as a potential approach for converting the “cold TME” phenotype of the persistent reservoir to a “hot TME” phenotype to enable immune clearance. Transforming “cold” TMEs to “hot” TMEs to enable clearance through antitumor immune activity has been an important cancer immunotherapy approach [36]. Further, inhibition of the top gene drivers of the TME score in the persistent reservoir may prevent transcriptional and signal transduction changes at the molecular level that drive the TME-like phenotype. Our analyses also revealed meaningful insights on short-lived, transient reservoirs. These reservoirs are effectively cleared by immune responses, providing a blueprint for determinants of curable HIV/SIV reservoirs. Determinants associated with these transient reservoirs highlight microenvironmental states that can be therapeutically targeted towards HIV cure approaches.

In summary, this study provided key insights on the hallmarks that distinguish persistent from transient HIV/SIV reservoirs, revealing the presence of several features consistent with those of tumor microenvironments **(Supplementary Table 1)**. In-depth spatial analysis revealed that at the transcriptional, local tissue neighborhood, cell, and host transcriptome gene levels, persistent reservoirs tend to reflect “cold”, immunosuppressed TMEs while transient reservoirs broadly reflect “hot”, immune-active TMEs. The identification of specific aspects of the VME that are shared with the TME provides key insights towards the effective application of immunotherapies for the eradication of one of the greatest barriers to a functional cure for HIV: persistent tissue reservoirs.

## Methods

### ImmunoPET/CT-guided spatial transcriptomics with complementary IF imaging

ImmunoPET/CT-guided necropsy was used as previously described to select tissue samples from SIVmac239-infected rhesus macaques for spatial transcriptomics analysis via the standard 10x Visium spatial transcriptomics platform. Spatial transcriptomics data were integrated with immunofluorescence imaging data as previously described [8].

### Ortholog conversion of rhesus macaque genes to human genes

Rhesus macaque to human orthologs were converted as previously described [8]. In brief, to convert rhesus macaque gene names to human orthologs, all gene names present in the spatial transcriptomics data were screened with a dictionary of rhesus macaque/human gene orthologs generated using the biomarRt package in R [86]. Upon determining which spatial transcriptomics genes had orthologs in the dictionary, at indices where an orthologous human gene name was available, the rhesus macaque gene name was replaced, in place at the corresponding index, to the human ortholog. Genes that mapped to multiple orthologs were not converted to prevent duplicate gene names.

### Spatial enrichment

Spatial enrichment analysis was completed using the decoupleR package (v2.1.2) in Python (v3.10.19) [35]. Raw data was read into Python and concatenated within each experimental condition (persistent reservoir vs. transient reservoir) and normalized using standard scanpy functions (scanpy.preprocessing.normalize, scanpy.preprocessing.log1p) [87]. Labels for SIV region (SIV-negative, SIV-neighbor, SIV-positive) were added to the metadata by spatial barcode. Gene sets for annotated diseases from the enrichDO package in R were loaded into Python and scored for spatial enrichment using the decoupler.pp.get_obsm function to obtain a composite score for each disease ontology condition at each spatial transcriptomics spot [34]. Several t tests were then completed with FDR correction, stratifying by SIV region, to determine which gene collections associated with various diseases were spatially enriched among each of the SIV groups in comparison to all other spots on each tissue. The function decoupler.tools.rankby_group was used to generate a data frame of spatial enrichments by SIV category.

Spatial enrichment overlays were visualized in Seurat [88]. Gene pathways from the enrichDO database were scored for the harmony integrated dataset combining the persistent and transient reservoir data for individual slide visualization using the scanpy.tl.score_genes command in Python [34, 89]. Cancer-associated gene scores that were negative were rounded to zero and the cancer gene pathway scores were visualized separately from and overlayed with the SIV density values at each spot. As the range of SIV density values in the persistent reservoir was smaller than the transient reservoir, the maximum range of SIV density that is visualized is up to the 98^th^ percentile of the score distribution of the integrated transient and persistent reservoir combined dataset while the cancer gene score distribution is up to the 90^th^ percentile (full SIV density distribution is shown in **Supplementary Figure 1**).

### ESTIMATE TME scoring

The tidyestimate package (v 1.1.1.9000) in R (v4.4.0) was used to complete ESTIMATE, immune, and stromal scoring for each spot on all spatial transcriptomics slides (**Supplementary Figure 2**) and for the full dataset after pseudobulking (using the AggregateExpression function in R) by persistent vs. transient reservoirs and SIV-negative vs. SIV-neighbor vs. SIV-positive regions [38]. Rhesus macaque gene names were converted to orthologs using the gprofiler package in R when applicable [90]. ESTIMATE, stromal, and immune scores were then visualized for each spot on each slide using Seurat [88]. Mixed linear effects modeling was completed using the lme4 and lmerTest packages in R to determine the statistical significance of group comparisons (i.e., persistent vs. transient, SIV-positive vs. SIV-neighbor vs. SIV-negative) [91].

### Neighborhood analysis

Neighborhood analysis was completed using the SpottedPy package (v0.1.10) in Python (v3.10.18) [39]. Raw spatial transcriptomics data were read into Python, concatenated into one anndata object, and gene names were converted to human orthologs where applicable. All 320+ of the gene pathway lists from the ImmunoOncology Biological Research (IOBR) database (v0.99.0) were then imported into Python from R (v4.4.0) [40]. IOBR gene pathways were scored only if the IOBR pathway contained at least one overlapping gene with the spatial transcriptomics data. Pathway scoring was completed using the scanpy.tl.score_genes function for the IOBR gene pathways. SIV density values at each spot were added to the metadata of the Python anndata object and SIV density scores less than 0 were rounded to 0. Scores of SIV density and TME pathways for each spot were then scaled to a range of 0 to 1 using the MinMaxScaler function from sckikit-learn [45]. “SIV-positive, high-density regions” (“hotspots” in SpottedPy) were then determined for each spatial transcriptomics slide with the spottedpy.create_hotspots function, using the additional arguments filter_columns= “SIV” and filter_value=1 to focus on regions of interest (ROI) containing SIV foci of infection. Hotspot distances for all spots within each slide were calculated using the spottedpy.calculateDistances function. Mixed linear effects statistical modeling was completed using the lme4 and lmerTest packages in R to determine which IOBR pathways were significantly closer in proximity to SIV-positive, high-density regions vs. SIV-positive, low-density regions by effect size (coefficient from mixed linear effects model * -log10(adjusted p value)) [92]. For the top 5 gene pathways in closer proximity to SIV-positive high-density regions or SIV-positive, low-density regions, mixed linear effects modeling was used to compare pathway scores within neighborhoods as well. Top pathways of interest were visualized and overlaid with SIV density neighborhoods using Squidpy (Spatial Single Cell Analysis in Python) standard visualization functions [93].

### Cell-cell interaction analysis

Cell-cell interactions were determined using the SOAPy (A package for Spatial-Omics in Python) package (v1.0.1) [41]. Raw spatial transcriptomics data were read into Python (v3.9.23) and concatenated into one anndata object. Ortholog conversion was completed for macaque to human genes as previously described [8]. Inferred cell type labels acquired through previously described methods were added to the metadata of the anndata object [8]. The label used for each spot was the maximum cell type frequency label merged with the SIV region category. Raw data were then preprocessed using standard scanpy functions (scanpy.preprocessing.normalize_per_cell and scanpy.preprocessing.log1p). Ligand-receptor pair data was acquired from the built-in SOAPy database using the soapy.tl.lr_pairs function, with the “human” database setting. Cell-cell interactions were then determined using the soapy.tl.cell_type_level_communication function to determine interactions at both the contact (local) and secretory (long-range) levels, using 1000 iterations. A secretory radius of 100μm was set for the analysis as the 10x Visium spatial transcriptomics used in the study contains spots that are 55μm in diameter, and therefore, spots within a 100μm radius are expected to be in direct contact. Raw cell-cell contact and secretory data were then extracted from the resulting anndata object and visualized using igraph in R to generate network plots of cell-cell interactions [94]. Network plots included the top 45% of cell-cell interactions per condition. The igraph degree function was used in R to determine the degrees of centrality of each cell type per condition. The pheatmap package was used to generate heatmaps of all contact and secretory cell-cell interactions within the persistent and transient reservoirs as well as the degrees of centrality of the cell types among the top 45% of interactions.

### Gene driver identification

(i) Gene list identification - Top gene drivers were identified through the generation of machine learning regression models. All data layers were joined for each condition (persistent vs. transient reservoir) using the JoinLayers function in R (v4.4.0). The raw data matrix with gene expression counts for each spatial transcriptomics barcode was extracted using the data@assays$Spatial$counts function in R as an input for machine learning regression. The ESTIMATE score (as described in the “ESTIMATE TME scoring” section) was the output variable for each barcode. Base models without tuning were created using the scikit-learn (v1.5.2) machine learning package in Python (v3.12.10) to test ten machine learning regression methods: Lasso Regression, Ridge Regression, ElasticNet Regression, Support Vector Regression, Decision Tree Regression, Random Forest Regression, K-Nearest Neighbors Regression, Neural Network, Gradient Boosting Regression, and Extreme Gradient Boosting Regression [45]. Accuracy of these initial baseline models was compared to determine the model with the highest accuracy of prediction of the ESTIMATE TME score, determined based on minimizing the mean squared error and maximizing the r^2^ of the model, resulting in the selection of Extreme Gradient Boosting (XGBoost) Regression. The optimal XGBoost model was then used to identify the top gene drivers. The optimized XGBoost model was run by generating a model in scikit-learn using a 75/25 test/train split, initiating an XGBoost Regressor using the xgb.XGBRegressor function from the xgboost package in Python, and selecting the top 1000 features from that initial model using the SelectFromModel and selector.fit_transform functions in scikit-learn. After selection of the top 1000 features, hyperparameter tuning was completed by doing a grid search using the GridSearchCV function to identify and rank the top features by importance by minimizing the mean rmse and maximizing the r^2^. The hyperparameter grid used for tuning spanned the following ranges: [max_depth: 3,5,7; min_child_weight: 1,5,10; learning_rate: 0.1, 0.2, 0.3; n_estimators: 100,250,500,750, 1000; gamma: 0,2,5; subsample: 0.5, 0.8, 1; lambda: 0,0.1,1; tree_method: hist].

(ii) Gene pathway enrichment among top gene drivers - Top gene driver enrichment was determined by running the enrichKEGG function on the unique gene drivers among the top 100 genes (for the persistent vs. transient reservoirs) ranked by importance, which was obtained from the machine learning regression modeling [95].

(iii) Top gene driver expression correlation with ESTIMATE scores - The spearman correlation between the gene expression (in the Seurat object SCT data layer) and the ESTIMATE scores at all barcodes was computed using the “cor” function in R. Negative correlations indicate an association of higher gene expression with a more TME-like phenotype because lower ESTIMATE scores are associated with the TME.

### Comparison to publicly available data

Upon identifying the top gene drivers using the methods described in the “Gene driver identification” section, each of the top five genes was queried in the National Cancer Institute’s GDC Data Portal to see which subtypes of cancers in the database tended to have high frequencies of mutations in the top gene drivers. To understand gene expression trends in the context of uterine corpus endometrial carcinoma, publicly available bulk RNAsequencing data from Luo et.al. [58] was visualized (using Enhanced Volcano [v1.24.0] in R [v4.4.2]), with annotations of genes that were present among the top drivers in the persistent reservoir [58].

## Acknowledgements

This research was supported in part through the computational resources and staff contributions provided by the Quest high-performance computing facility and Sequencing Core facility (NUSeq) at Northwestern University, which is jointly supported by the Office of the Provost, the Office for Research, and Northwestern University Information Technology. This research was also supported in part through the computational resources and staff contributions provided by the Genomics Compute Cluster, which is jointly supported by the Feinberg School of Medicine, the Center for Genetic Medicine, and Feinberg’s Department of Biochemistry and Molecular Genetics, the Office of the Provost, the Office for Research, and Northwestern Information Technology. The Genomics Compute Cluster is part of Quest, Northwestern University’s high-performance computing facility, with the purpose of advancing research in genomics.

## Funding sources

This research was supported by NIH/NIAID funding for HIV research (P01AI169600 and R01MH125778 to T.J.H and R.L-R; R37AI094595, 1U54AI170856-01, and R01AI177265 to T.J.H.; P01AI131346 to T.J.H.), and NIH/NIAID funding for the Third Coast Center for AIDS Research (P30 AI117943 to R.L-R).

## Data Availability

All scripts and bioinformatic pipelines are available at the Lorenzo-Redondo lab github (https://github.com/rlr-lab/). All processed files and imaging files are stored on the Northwestern FSMResFiles centralized data storage platform for long-term storage. Sequencing files are deposited to NCBI as Bioproject.

## Author contributions

EUC: Conceptualization, Data curation, Formal Analysis, Investigation, Methodology, Visualization, Writing – original draft, Writing – review & editing. NS: Visualization, Writing – review & editing; AM: Conceptualization, Writing – review & editing; TJH: Conceptualization, Funding acquisition, Investigation, Project administration, Resources, Supervision, Validation, Writing – original draft, Writing – review & editing; RL-R: Conceptualization, Data curation, Formal Analysis, Funding acquisition, Investigation, Methodology, Project administration, Resources, Supervision, Validation, Visualization, Writing – original draft, Writing – review & editing. All the authors read, reviewed, and approved the final manuscript.

## Notes

### Competing Interest Statement

The authors have declared no competing interest.

