## Supplementary Table 1 for "A Tissue Microenvironment Analogous to Certain Tumor Microenvironments Facilitates HIV Persistence"

**Supplementary Table 1:** Characteristics of the Persistent versus Transient Reservoir that Reflect Aspects of the Tumor Microenvironment

| | **Persistent Reservoir** | **Transient Reservoir** |
|---|---|---|
| **Cancer gene scoring spatial distribution** | Localized enrichment of cancer-associated genes in SIV-associated regions, with surrounding collagen/fibrotic-related processes | Broad enrichment of cancer-associated genes with relatively higher enrichment in SIV-associated regions |
| **Immune infiltration** | ↑ immune infiltration in SIV-associated regions | ↑↑↑↑↑ immune infiltration in SIV-associated regions |
| **Gene Pathways** | SIV-positive, high-density areas: energetic metabolic pathways<br><br>SIV-positive, low-density areas: stromal activation, EMT, TGFβ, TNF, and CAF-associated pathways | SIV-positive, high-density areas: Activation of $CD8^+$ T cell, antigenic, and nucleic/amino acid metabolic pathways |
| **Cell Interactions** | Tregs receive signals from various, notably innate, immune cells in the microenvironment | Tregs send signals to various adaptive immune cells in the microenvironment<br><br>Contact interactions of SIV-associated Th17 and $CD8^+$ T cells |
| **Top Unique Host transcriptional markers** | KRT8<br>EPCAM<br>MLEC | KRT18<br>TSPAN8<br>SASH3 |
